# Off-target metagenomics: Leveraging whole genome sequencing to study the bacteriome of the liverwort *Calasterella californica*

**DOI:** 10.1101/2025.01.23.634585

**Authors:** Ixchel González-Ramírez, Michael J. Song, Elijah C. Mehlferber, Brent D. Mishler

## Abstract

**Premise of study:** The recovery of non-target organism reads, especially when whole organisms are sampled, constitutes a great opportunity for studying microbial communities. The increase in whole genome sequencing feasibility, and the development of new pipelines and databases enable the use short reads to study bacterial communities associated with organisms.

**Methods:** We utilized population genomic data of the liverwort *Calasterella californica* obtained through the California Conservation Genomics Project to characterize the composition of its associated bacterial communities and explore its variation across the geographic space.

**Key results:** The bacterial communities associated with *C. californica* were dominated by the methanotroph *Methylobacterium* and other Hyphomicrobiales, a group that includes well known plant symbionts. While diversity metrics of bacteria composition was similar across localities, we found significant differences in the relative abundance of a few taxa across California regions, likely driven by differences in precipitation and temperature seasonality.

**Conclusions:** Our results support previous observations that liverwort bacterial communities are not randomly assembled, suggesting a potential role of the plant in determining community composition, an emerging pattern that deserves more attention. Our novel off-target metagenomics approach can be applied to any population level re-sequencing where whole organisms are sequenced, opening the door to exciting avenues of microbiome research using re-purposed data from landscape genomics.

## Introduction

An important opportunity for leveraging existing sequencing data lies in the recovery of non-host reads that are inadvertently captured alongside the target organism during whole-genome sequencing (WGS). The standard protocols used for WGS extraction do not distinguish between the DNA of the target organism and other organisms that might be present in the sample—such as parasitic or symbiont microorganisms—which are often considered contaminants (Steinegger and Salzberg 2020). In larger hosts, whose extractions are usually performed on samples of internal organ-tissue (Wood et al. 2022; Benham et al. 2023; Supple et al. 2024), we might expect a relatively low proportion of contaminant reads. But performing WGS on small organisms, like insects, small invertebrates, and plants such as bryophytes, often involves using complete individual(s) as a sample, which increases the proportion of non-host reads that we might expect because of the initial proportion of host tissue is smaller, and also because sequencing whole organisms includes environment-interfacing tissues. Given the growing appreciation for studying holobionts—*i*.*e*., an organism and all the associated organism living within and around it (Vandenkoornhuyse et al. 2015)—we can leverage WGS data of whole organisms to provide insights into bacterial communities, symbiosis, and co-evolution between host and micro-organism (Song et al. 2025a, 2024), in an approach that we refer to as “off-target metagenomics”.

To demonstrate the practicality of the off-target metagenomics approach, we re-purpose data generated by the California Conservation Genomics Project (CCGP), originally intended to create genomic maps across the state to influence land management and conservation policy (Shaffer et al. 2022; Toffelmier et al. 2022; Fiedler et al. 2022; Beninde et al. 2022), to explore the microbiome associated with a liverwort species. The CCGP collected and sequenced samples for hundreds of individuals of more than 100 species across California. Many of these focal taxa represent an ideal opportunity to analyze the non-target reads to study the composition of microbial communities and their potential variation across the geographic space, especially for those taxa where organisms were sequenced whole (Mead et al. 2024; Grether et al. 2023; Blair et al. 2022; Huang et al. 2022; González-Ramírez et al. 2025a; Song et al. 2025b). Offtarget metagenomics is possible thanks to the recent development of a metagenomic profiling software, MetaPhlan4, (Blanco-Míguez et al. 2023) that matches short-reads from metagenomic samples to “species level genome bins” (SGBs). Using SGBs as matching bins improves the profiling of undescribed microbial communities in comparison to other approaches like reference-based computational ones—which are limited by the number of reference genomes available—or metagenome-assembled genomes (MAGs)—which require high coverage for all the taxa (Blanco-Míguez et al. 2023). Together, population level datasets, and analytical tools like MetaPhlan, allow for research into microbiome geographic variation which has hitherto been greatly overlooked (Härer and Rennison 2023), especially for understudied taxa like liverworts.

In this study, we focused on the bacterial community of the liverwort *Calasterella californica* (Fig. 1, González-Ramírez et al. (2025a)). Liverworts are small plants with comparatively little tissue differentiation and many cells in direct contact with the environment. Like many plants, liverworts are known to have important associations with microorganisms, particularly fungi. *C. californica* has a geographical distribution, occurring in contrasting environments that range from wet redwood forests to the dry Anza-Borrego desert in Southern California (González-Ramírez et al. 2025b), making it an ideal system to investigate the variation of its associated bacterial community across the geographic space. In recent years, there has been an effort to understand the microbial communities associated with liverworts (*e*.*g*., Wicaksono et al. 2023; Alcaraz et al. 2018; Kutschera and Koopmann 2005; Young et al. 2025) that point to potentially stronger associations than previously thought, and highlight the need for further investigation. By applying the off-target metagenomics approach to *C. californica* we provide new evidence on the potential tight relationships of liverworts and their bacteriomes, and address the question of how much these bacterial communities vary across the geographic space.

**Figure 1:**
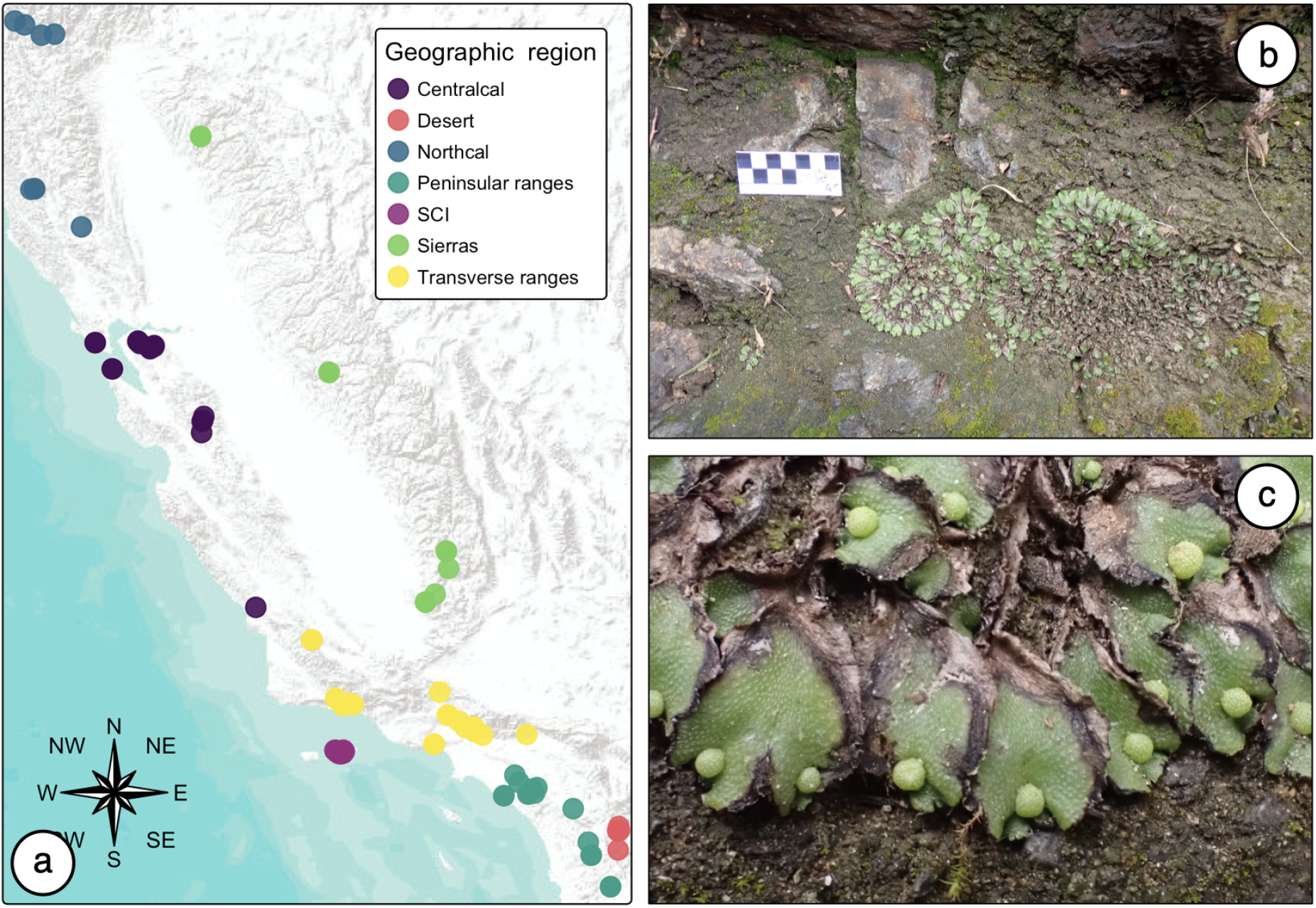
**a**. Collection locations of *Calasterella californica* samples for this study, dots are color coded by their California geographic region. **b**. Typical morphology, *C. californica* grows on bare and exposed soil or rocks, forming circular mats. **c**. Close up to female thalli of *C. californica* displaying the proximity of this plant with the soil.

## Methods

### Sampling

*—*The original sampling of *C. californica* consisted of 110 samples collected as part of the CCGP project (González-Ramírez et al. 2025a). This sampling focused on representing the broad geographic space and environmental conditions in which this liverwort occurs. The localities where the samples were collected were assigned to different California geographic regions that largely reflect shared environmental conditions and shared geological history (Individual samples geographic regions are recorded in Sup. Tab. S.2). The Northern California region is characterized by higher precipitation and cooler temperatures. The Sierras region spans the interior mountain range of California, east of the Central Valley and it is characterized by a shared geological history, marked seasonal variation, and higher altitude. Central California samples span the coastal region (west of the Central Valley) from the Bay Area to the Transverse Ranges. This region is characterized by coastal scrublands (as opposed to more perennial conifer forests in Northern California), and comparatively less seasonal variation than samples from more inland regions. In Southern California, we differentiate between the Transverse Ranges, characterized by higher altitude, precipitation, and seasonality than the more coastal Peninsular Ranges. The Santa Cruz Island region is very similar to the Central California region in terms of vegetation, but the strong oceanic influence decreases the seasonal variation in temperature and rainfall. Finally, the desert region corresponds to the flat area east of the Transverse Ranges that belongs to the Sonoran desert and is characterized by a dry and warm climate (Fig. 1 and Sup. Fig. S2).

We leveraged 97 of the CCGP samples to characterize *C. californica*’s bacteriome across the geographic space. Only 97 were used out of 110 liverwort collections because the remaining 13 samples did not pass the sequencing quality checks or were found to be a different liverwort species. Each liverwort sample was collected in a different geographic location. At each location, one to two clumps of liverworts were collected in hard plastic containers and transported to the lab. From each collection, the largest 1-3 thalli were isolated and cleaned by manually removing soil particles and most of the rhizoids and used for DNA extraction. The number of thalli used was based on securing similar amount of tissue across samples. When more than one thallus was used, we selected thalli that, according to the growth pattern, appeared to be part of the same individual. Previous microbiome studies have shown that short term storage conditions have a surprisingly small impact on bacterial diversity and community structure as long as the storage strategies are consistent across samples and that care is taken to limit the amount of freeze-thaw cycles (reviewed in Goodrich et al. (2014)).

### DNA-extraction, sequencing, and quality checks

*—*For each sample, total genomic and symbiont DNA of one to four thalli was extracted using either the Qiagen DNeasy Plant Pro kit (California, USA) or a CTAB protocol. Library preparation and sequencing was carried out at the UC Davis DNA Technologies & Expression Analysis Core Laboratory. Libraries were sequenced using an Illumina NovaSeqX 300 (PE150) flow-cell. During ealy sequencing quality checks, all the samples displayed a bimodal GC-content distribution that characterizes the presence of both bacterial (high GC-content), and liverwort (low GC-content) DNA.

### Reads processing and bacteriome characterization using MetaPhlAn

All the paired-end files were quality checked and cleaned from adaptors using `Trimmommatic` (Bolger et al. 2014). For each sample, the paired-end reads were mapped to the *C. californica* IGR150-G1 reference assembly (González-Ramírez et al. 2025a) using `bwa-mem` (Li 2013). Next, using `‘samtools’` (Li et al. 2009) with the flag -f 12, we kept only the reads whose mate was also unmapped to the liverwort reference genome, and these reads were used as the input to study the microbiome. To characterize the *C. californica* metagenome composition of each liverwort, we used `MetaPhlAn,` a recently developed pipeline that uses a compendium of metagenome-assembled genomes and reference genomes of microbes to characterize microbial communities and estimate relative abundances from shotgun sequence data (Blanco-Míguez et al. 2023). This approach differs from other common microbiome-targeting approaches such as 16S rRNA/18S rRNA/ITS sequencing or the assembly and annotation of full metagenome assembled genomes (MAGs) in that it does not attempt to assemble the microbial markers or genomes, but instead directly uses the short-reads to characterize the metagenome. We ran MetaPhlan on the Berkeley Computer Cluster SAVIO using the most recent reference database of SGBs provided by the MetaPhlan team of developers (*mpa_vJan*25_*CHOCOPhl AnSGB_*202503), and employing 20 cores simultaneously for each sample. The number of reads a each step of this workflow is detailed in Sup. Tab. S.2.

### Bacteriome composition and diversity

*—*All the statistical analyses and visualizations were performed using the `microeco` package (Liu et al. 2025) in R (R Core Team 2000). The output obtained from MetaPhlan was formatted as input for microeco using the R package `file2meco` (Liu et al. 2022). To quantify the *α*-diversity for each sample we used multiple standard diversity indexes: Shannon (1948), Simpson (1949), Pielou (1966), and coverage, as implemented in `microeco`. We tested the potential effect of geographic region on these diversity metrics using an Analysis of Variance (ANOVA) (Fisher 1928).

To characterize the similarity among the bacteriomes of the different samples of *C. californica*, we calculated a Bray–Curtis dissimilarity matrix and computed a Non-Metric multidimensional scaling (NMDS) ordination. To assess whether samples belonging to the same geographic region were more similar in composition we performed a PerMANOVA, as implemented in `microeco` (Anderson 2001). Furthermore, we evaluated whether or not specific taxa were significantly over or under represented across geographic regions using a differential abundance test at the family and genus taxonomic levels. We assessed significance using a linear discriminant analysis effect size (LefSE) as implemented in `microeco` (Segata et al. 2011). Before using this test, we applied an arc sine square root transformation of the abundances, to account for the use of relative abundances.

### Effect of climatic variables on microbial community structure

*—*Since macro-environmental variables might affect the bacteriome of *C. californica* by determining the pool of bacteria available in the environment, and because this liverwort occurs in a wide range of climatic conditions, we evaluated the effect of climatic variables on the composition of *C. californica*’s bacteriome. For this, we extracted the values of the bioclimatic layers from the BioClim database (Fick and Hijmans 2017) for each collection point using the R packages sf (Pebesma 2018) and raster (Hijmans et al. 2013). Of the 19 variables in BioClim we selected a subset of variables with relatively low correlation and high variation within our geographic scope (Sup. Fig. S1. We used a Correspondence Analysis (CCA) to visualize the relationship between the climatic variables and the microbial composition of *C. californica* samples at the taxonomic family level (Ter Braak 1986), and we tested for the significance of this relationship, using a Mantel test implemented in `microeco` (Mantel 1967).

### Functional composition

*—*In order to characterize the functional profile of the bacteriome associated to *C. californica*, we used the `FAPROTAX v1.2.10.` database (Louca et al. 2016a,b) to link the taxonomic composition of the bacteriome of each sample to functional diversity using the `microeco` built in function. Finally, we tested for differential abundance of these functions in different geographic regions using a linear discriminant analysis effect size (LefSE) with the relative abundance of the different functions in the bacteriome as input.

The bash, python, and R code used to perform the cleaning, microbiome pipeline and statistical analyses and visualizations is available in the Github repository: https://github.com/ixchelgzlzr/CCal_microbiome.

## Results

### Bioinformatics

Ninety-seven of the original 110 *C. californica* passed quality control and were used in our study (Sup. Tab. S.2). The number of sequences before filtering steps were on average around 30,600,000 reads. The smallest sample contained 13,028,666 reads and the largest contained 45,988,969 reads. After filtering out host *C. californica* reads, we were left with around 19,000,000 reads on average (61% of the total), with the smallest sample containing 3,835,559 reads and the largest 36,016,561 reads. When we profiled the microbiome using MetaPhlan, on average 96.63 percent of reads were unclassified and not used for downstream analysis. Our final analysis on classified microbiome reads is composed of on average around 640,000 reads per sample with the smallest sample containing 70,519 reads and the largest 3,610,608 reads (Sup. Tab. S.2).

For our dataset, the running time on a standard laptop using eight cores was about seven hours per sample. In our particular cluster environment (SAVIO, at UC Berkeley), we observed that even with 384 GB of total RAM available for 20 cores, we faced memory constraints when trying to parallelize per sample. We found that we obtained the best performance when assigning all the cores and RAM memory to a single sample, and running the analysis sequentially for all samples. With this set up, each sample took about 30 min (20 cores simultaneously with 384 GB of RAM).

### Composition of the bacterial microbiome of C. californica

Across the 97 samples of *C. californica*, the MetaPhlan pipeline identified 751 species level bacterial OTUs, belonging to 338 different genera, 141 families, 87 orders, 51 classes, and 15 phyla (Sup. Tab. S.3). Overall, the bacterial microbiome of *C. californica* is dominated by bacteria in the order Hypomicrobioales, often accompanied by other bacteria orders like Pseudomononadales, Pseudonocardiales, Burkholderiales, Mycobacteriales, Micrococcales, and Oscillatoriales (Fig. 2).

**Figure 2:**
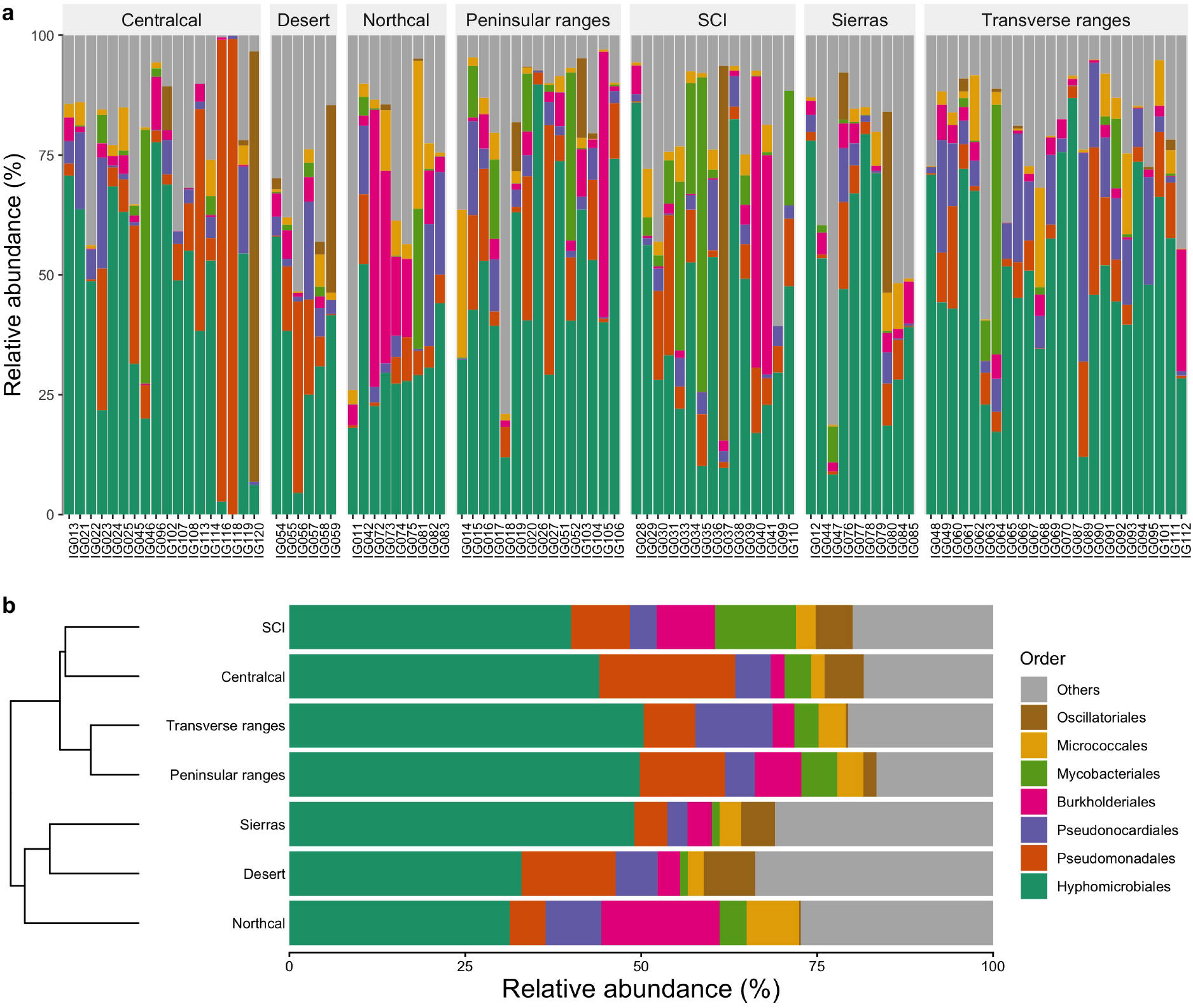
Composition of the bacteriome of *C. californica*. **a**. Relative abundance of the seven most abundant orders of bacteria across all the samples of *C. californica*. The samples are arranged by the geographic region where they were collected. **b**. Composition grouped by geographic region. The relative abundances are obtained from group averages. The dendrogram on the left reflects similarity among regions based on a hierarchical clustering calculated from Euclidian distances.

An ANOVA test did not find any of these *α*-diversity metrics to be significantly different between geographic regions (Sup. Tab. S.1). Nevertheless, a PERMANOVA test (*P <* 001, *R*^2^ = 0.127,*F* = 2.19 *d f* = 6) found that there is a weak but significant effect of the geographic region on explaining the patterns of *β*-diversity of the bacteriome of *C. californica* (Fig. 3). The linear discriminant effect size (LefSE) analysis revealed that there are 18 families and 50 genera of bacteria that have significantly differential abundance in different geographic regions (Fig. 4).

**Figure 3:**
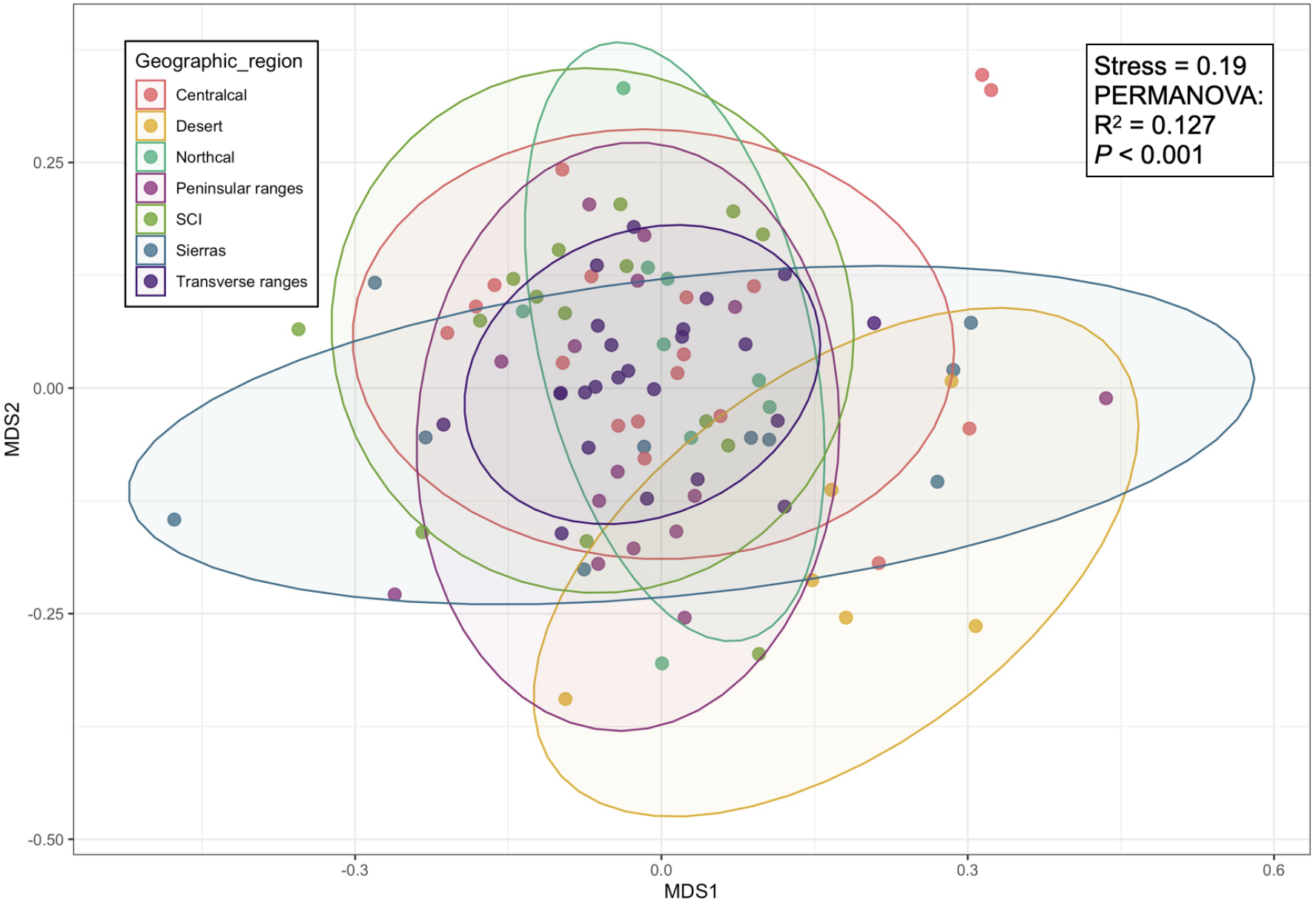
Non-metric multidimensional scaling (NMDS) ordination of *C. californica* associated bacterial communities based on Bray–Curtis dissimilarities. Each point represents a different sample color-coded by the geographic region it was collected. While the biplot ordination does not show a strong arrangement by geographic region, a PERMANOVA test finds a weak but significant effect of the region on the composition of the bacterial community.

**Figure 4:**
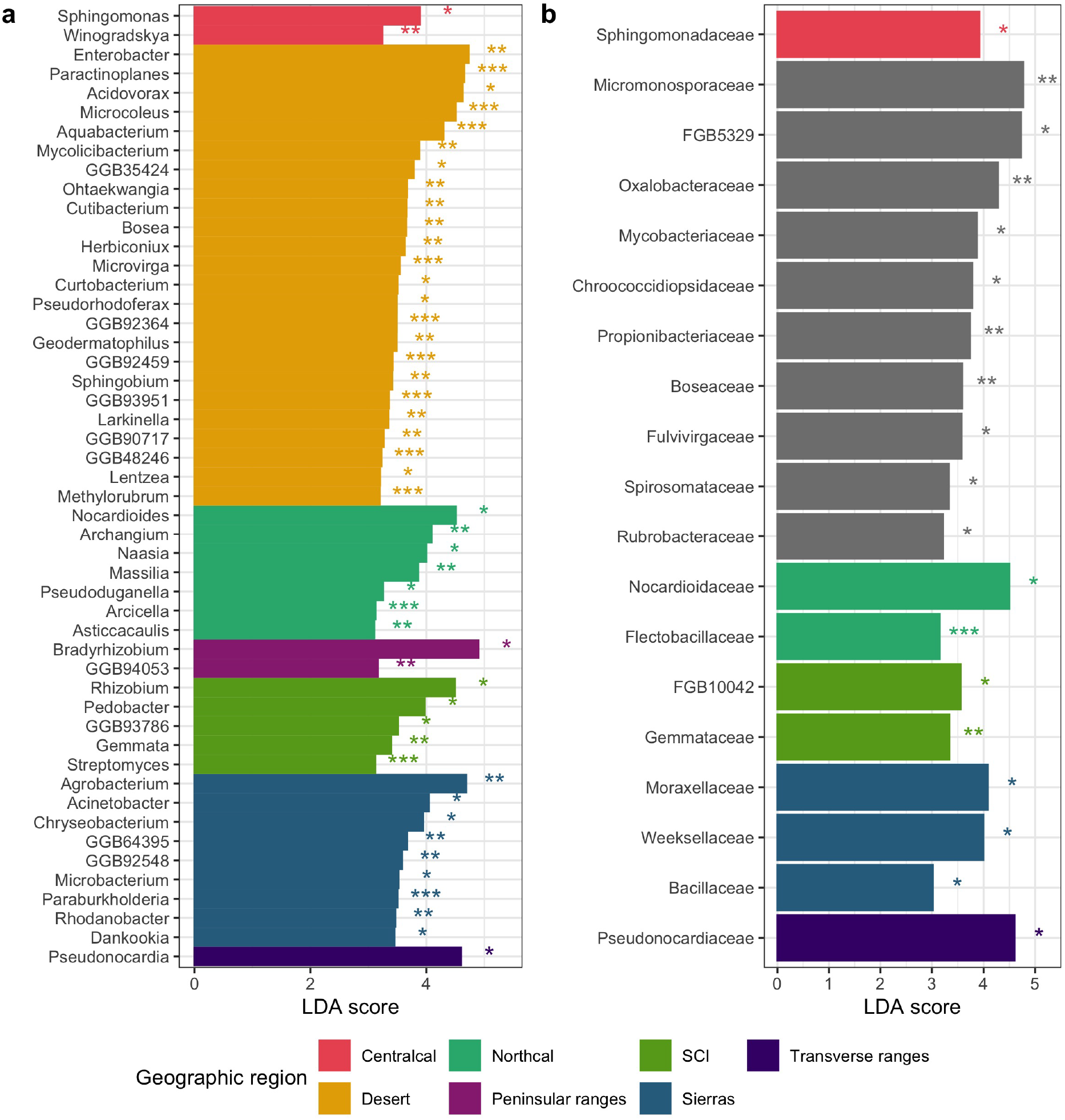
Significant differential abundance of bacterial (a) genera and (b) families across geographic regions. There are 18 families and 50 genera of bacteria with significantly different abundances across geographic regions. Differential abundance was determined based on LDA scores *>* 3. Significance values are signaled with asterisks such as *P <* 0.5*\**; *P <* 0.1*\*\*, P <* 0.01*\* * \**. When differential abundance is characteristic of a single region, the bar is colored by such region, otherwise the bar is grey. Grey bars imply that more than one region has differential abundance of this taxon.

### Effect of macro-climatic variables in bacteriome composition

The three macro-climatic variables that we selected to test for the effect of climate on bacteriome composition (via affecting the pool of bacteria in the environment) were temperature seasonality, altitude, and annual precipitation. These three variables explained most of the variation across geographic regions and captured the variation in other highly correlated variables (Fig. S2). A biplot visualization of the correspondence analysis shows an significant association of annual temperature with the main ordination axis according to a permutation analysis (*χ*^2^ = 0.17, *F* = 2.39, *P* = 0.002), which also correlates with the abundance of the taxa shown in black arrows: Oscillatoriaceae, Sanguibacteriaceae, Calotrichaceae, Aestuariivirgaceae, Coleofasciculaceae, Dinobryaceae, Perlucidibacaceae, Leadbetterellaceae, FGB26132, and FGB52975 (Fig. 5). And the second ordination axis has a marginal association with temperature seasonality (*χ*^2^ = 0.097, *F* = 1.35, *P* = 0.055; Fig. 5). Additionally, a Mantel test also supports a weak but significant effect of annual precipitation and temperature seasonality (but not altitude) in the composition of the bacterial communities of *C. californica* (Mantel test: temperature seasonality, altitude, annual precipitation; Pearson correlation coefficient: 0.11, 0.07, 0.18; adjusted P-values: 0.006, 0.064, 0.006, respectively).

**Figure 5:**
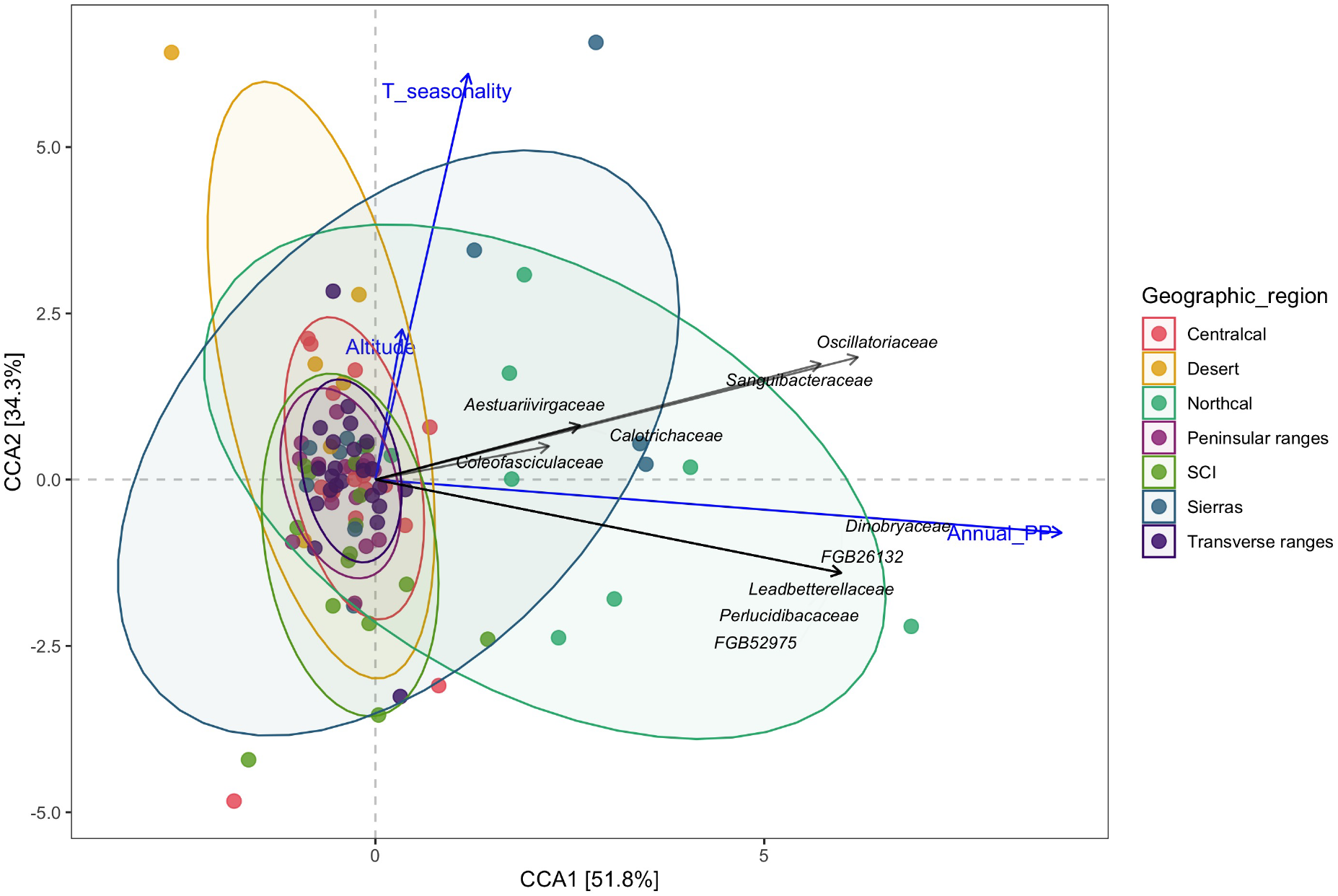
Correspondence Analysis (CCA) of the bacterial communities of *C. californica*. The blue arrows show the directionality of the variation of climatic variables in the multivariate space. The black arrows show the main axis of variation of bacterial families.

### Functional composition

The functional composition analyses shows that there is a relatively high proportion of bacteria associated with energy sourcing, specifically performing the functions of aerobic chemoheterotrophy and anaerobic chemoheterotrophy. Other functions that are relatively well represented in the bacteriome of *C. californica* are methylotrophy, methanotrophy, hydrocharbon degradation, ureolysis and methanol oxidation (Fig. 6). All of these highly represented functions seem to be common across all the samples. A differential abundance LefSE analysis performed for these different functions showed significant differences in 20 categories (Fig. S4). However, none of these taxa in any category comprised more than four percent of the total relative abundance (Fig. S3).

**Figure 6:**
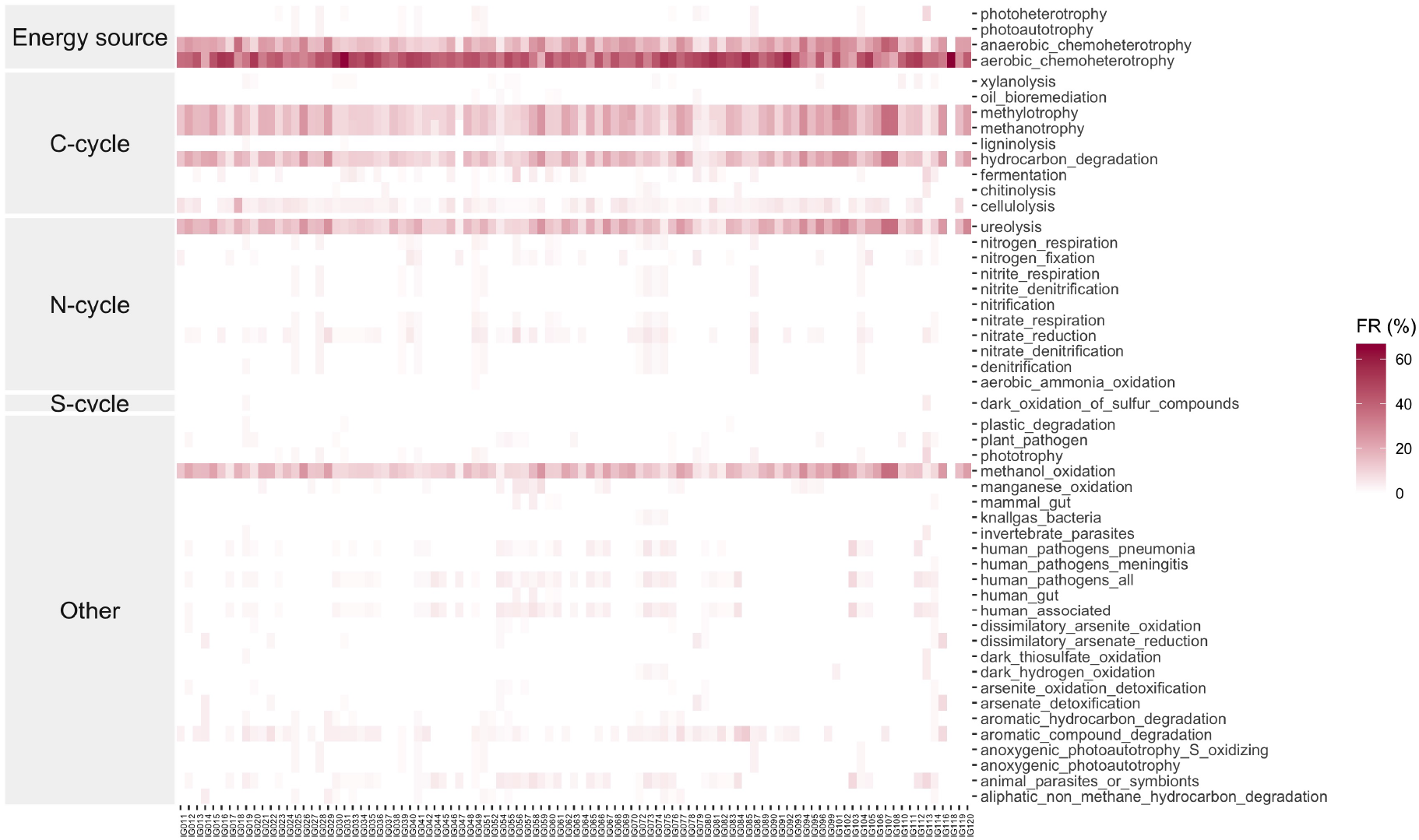
Functional composition of the bacteriome of *C. californica*. Relative abundance of bacterial functions in the community of bacteria associated to 97 samples of *C. californica*.

## Discussion

Despite living in contrasting regions across California (Fig. 1 and Fig. S2), the bacterial communities associated with *C. californica* are similar in their *α*-diversity metrics across all geographic regions (Tab. S.1). This overall similarity extends to their composition—*i*.*e*., all bacterial communities are dominated by bacteria in the order Hyphomicrobiales, with important contributions from a few other orders, such as Pseudomononadales, Pseudonocardiales, Burkholderiales, Mycobacteriales, Micrococcales, and Oscillatoriales (Fig. 2). These results are congruent with previous work in liverworts of the genus *Riccia*, whose bacterial communities are not affected by environmental conditions. For example, Wicaksono et al. (2023), using a traditional targeted 16s sequencing approach, found a stable bacterial microbiome in *Riccia* individuals growing in contrasting soil types. Similarly, Wiśniewski et al. (2025), using nanopore long read sequencing, identified a core microbiome, that was significantly enriched on some bacterial taxa in comparison to surrounding soil. While comparative studies on the bacterial communities associated with liverworts are still scarce, these three studies together provide growing evidence that liverwort-associated bacterial communities are non-random and stable regardless of the environmental conditions in which individual liverworts occur. This is in contrast with patterns observed in angiosperms, where bacterial community composition is strongly shaped by both host genotype and environment (Wagner et al. 2016; Wei and Tan 2023; He et al. 2024).

Among the taxa that were consistently abundant in *C. californica* bacterial communities were bacteria in the order Hyphomicrobiales. Two representatives of this order, *Methylobacterium* and *Rhizobium*, were also found to be main components of the microbiome of the model liverwort *Marchantia polymorpha* (Alcaraz et al. 2018), where they play an important role promoting the growth of this plant (Kutschera and Koopmann 2005). Hyphomicrobiales contain many genera that are known to be beneficial to plants not only for stimulating plant growth, but also as nitrogen-fixers and root nodulation promoters (Lindström and Mousavi 2020). The functional profiles of the bacteriomes were also very conserved across samples and geographic regions (Fig. 6) with high relative abundance of bacteria that perform methanol oxidation, methano- and methylotrophy, and aerobic chemoheterotrophy. These results highlight the role of that methylotrophic bacteria play as the dominant taxon in the *C. californica* microbiome. Whether and how liverworts actively maintain high proportion of these bacteria in their microbiome, in a dynamic and heterogeneous environment, warrants more investigation.

While the primary components of the microbiome of *C. californica* are the same across samples, when scrutinizing the identity of less abundant taxa, we found some differences across geographic regions (Fig. 4) potentially associated with variation in annual precipitation and temperature seasonality (Fig. 5). For example, higher abundance of photosynthetic bacteria in the families Oscillatoriaceae, Calotrichaceae, and Coleofasciculaceae was with higher annual precipitation. Desert samples had many significantly differentially abundant genera (Fig. 4), many of which—as reflected in the functional profile (Fig. S4)—are plant pathogens and human, animal, and gut associated (Fig. 4). One potential explanation is simply that the samples from the desert were collected near seasonal streams influenced by runoff and touristic human activities. On the other hand, previous work in angiosperms has documented shifts of bacteria composition associated with water availability (*e*.*g*., Chao et al. 2025) and disease response (*e*.*g*., Gao et al. 2021), suggesting some mechanisms that might be affecting these secondary bacterial components of *C. californica* microbiome. Further work would need to be done in order to understand the cause of relative abundance differences in minor components of *C. californica*’s microbiome in the most extreme sampled localities.

Using a novel off-target meta-genomics approach, we were able to characterize the bacterial composition of the *C. californica* microbiome. Although a large fraction of off-target (*i*.*e*., non-host related) reads were unclassified, this reflects MetaPhlan’s focus on bacterial metagenomes, while we expect to have reads associated with other groups such as fungi, small metazoans, and viruses. With adequate analytical tools, these non-bacterial reads represent exciting opportunities to investigate other organisms associated with this plant. A real limitation of this approach—which is shared among all metagenomic studies—is the incompletness of the reference databases, which are potentially underrepresenting the diversity of bacteria in uncommonly studied environments. This characteristic of our databases highlights the need for continuous work on building larger genomic datasets (whether they are MAGs, 16S, SGBs).

While our approach, by definition, leverages data that was not aimed to study bacterial communities, results obtained with this approach are complementary to other microbiome targeted approaches, and particularly valuable to identify questions that require more targeted research. Furthermore, the congruent results of our study with previous research on liverwort microbiome studies that used 16S rRNA metabarcoding (Wicaksono et al. 2023; Alcaraz et al. 2018) is not only interesting insofar that they imply general patterns of microbiome composition across liverworts, but also because they provide indirect support that 16S rRNA metabarcoding and metagenomic approaches like MetaPhlAn discover patterns that are consistent with each other. Additionally, shotgun sequencing, such as our study, has been found to have more power in identifying rare taxa than 16S (Durazzi et al. 2021), as it is not limited by primer amplification bias (Campanaro et al. 2018), and has greater taxonomic resolution than 16S (Laudadio et al. 2018).

Currently, there has been an increasing emphasis on incorporating long-read sequencing technology, which has facilitated the assembly of high quality metagenome assembled genomes and allowed for strain-level resolution and metabolic profiling (Han et al. 2024). Nonetheless, these approaches remain both cost-prohibitive and computationally expensive. At the same time, population genomic studies that use short-read shotgun resequencing approaches are becoming more common. The use of MetaPhlAn in our study exemplifies how sequencing produced for plant genomics work can be leveraged to obtain microbiome metagenomics information. Our results demonstrate that meaningful metagenomic insights can be made from utilizing these types of landscapes genomic datasets, and these insights are especially impactful for groups relatively understudied, as are liverworts. As the field moves toward long-read sequencing, this paper shines a spotlight to a potentially large body of microbiome research that can investigated that would otherwise be unused.

Our results are consistent with previous knowledge on liverworts bacterial microbiomes generated through standard methods to study microbiomes, pointing to the efficacy of these data-mining strategies to do exploratory studies that lead to targeted research questions on the assembly of microbial communities. In the case of liverwort bacteriomes, it is increasingly exciting to conduct sampling and *in vitro* studies aimed to understanding the likely active role that liverworts play in the assembly of its bacterial communities.

## Acknowledgements

The authors thank Carl Rothfels for his constructive comments and edits to the manuscript. Thank you to two anonymous reviewers and two editors for their thoughtful and constructive feedback of this manuscript. Thank you to the California Conservation Genomics Project, with funding provided to the University of California by the State of California, State Budget Act of 2019 [UC Award ID RSI-19-690224]. IGR was supported by a UC Mexus-CONACyT fellowship (number 709967), a Plant Science Fellowship by Oak Spring Garden Foundation, and the Philomathia Graduate Fellowship in Environmental Sciences, at UC Berkeley. This research used the Savio computational cluster resource provided by the Berkeley Research Computing program at the University of California, Berkeley (supported by the UC Berkeley Chancellor, Vice Chancellor for Research, and Chief Information Officer).

## Author contributions

IGR and MJS designed the study, performed the analyses and wrote the first draft of the manuscript. ECM contributed to analyses and edition of following versions of the manuscript. BDM supervised the work and edited the manuscript.

## Data availability statement

All the data and code used for the analyses is hosted in the GitHub repository:

https://github.com/ixchelgzlzr/CCal_microbiome.

